# ElasticBLAST: Accelerating Sequence Search via Cloud Computing

**DOI:** 10.1101/2023.01.04.522777

**Authors:** Christiam Camacho, Grzegorz M. Boratyn, Victor Joukov, Roberto Vera Alvarez, Thomas L. Madden

## Abstract

**Background:** Biomedical researchers use alignments produced by BLAST (Basic Local Alignment Search Tool) to categorize their query sequences. Producing such alignments is an essential bioinformatics task that is well suited for the cloud. The cloud can perform many calculations quickly as well as store and access large volumes of data. Bioinformaticians can also use it to collaborate with other researchers, sharing their results, datasets and even their pipelines on a common platform.

**Results:** We present ElasticBLAST, a cloud native application to perform BLAST alignments in the cloud. ElasticBLAST can handle anywhere from a few to many thousands of queries and run the searches on thousands of virtual CPUs (if desired), deleting resources when it is done. It uses cloud native tools for orchestration and can request discounted instances, lowering cloud costs for users. It is supported on Amazon Web Services and Google Cloud Platform. It can search BLAST databases that are user provided or from the National Center for Biotechnology Information.

**Conclusion:** We show that ElasticBLAST is a useful application that can efficiently perform BLAST searches for the user in the cloud, demonstrating that with two examples. At the same time, it hides much of the complexity of working in the cloud, lowering the threshold to move work to the cloud.

## BACKGROUND

BLAST (Basic Local Alignment Search Tool) [1] is used by biomedical researchers to characterize sequences by identifying similar sequences, with the command-line BLAST+ package [2] used for pipelines as well as tasks with large numbers of searches or custom databases. The BLAST+ package supports all types of possible searches (e.g., nucleotide-nucleotide, protein-protein, protein-translated nucleotide, profile searches etc.), user-provided databases, a built-in limit by organism feature, and multiple report choices. BLAST+ is supported on LINUX, Mac, and Windows.

The rapid growth of GenBank (see Table 1 in [3]) results in a continuing increase in the size of the most popular BLAST databases, requiring more effort to host them locally and more computational power to run searches. At the same time, cloud computing has become mature and offers an opportunity for Bioinformaticians [4]. It provides infrastructure such as instances, which are virtual servers that can be started on-demand and can contain different numbers of virtual CPUs (vCPUs) and different amounts of memory. It also provides object storage (cloud buckets) that can hold large amounts of data independent of any server as well as advanced tools to help orchestrate workflows. Cloud computing supports collaboration, with researchers able to easily share data, workflows, and compute environments with colleagues at other institutions. [4]. To enable cloud computing, the NCBI is now hosting popular BLAST databases on Amazon Web Servers (AWS) and Google Cloud Platform (GCP) [5], which users can easily download to their instance. The NCBI is also hosting 25.6 PB of SRA data in AWS and GCP (as of September 2021) [6]. The NIH Science and Technology Research Infrastructure for Discovery, Experimentation, and Sustainability (STRIDES) [7] initiative encourage the use of the Cloud by biomedical researchers. The cost model for the cloud (pay for usage) can present difficulties, but authors [4, 8, 9] discuss best practices to minimize costs. Multiple groups have demonstrated that the cloud is a viable platform for bioinformatics [4, 10, 11, 12].

**Table 1:** Search sets used for the taxonomic classification (derived from GTax database).

| Search set | Number of sequences | Number of bases |
| --- | --- | --- |
| Eudicotyledons | 880 | 39,174,504,447 |
| non-eudicotyledons | 41,218 | 447,368,164,093 |

Scheduling BLAST+ searches on the cloud involves several steps which include bringing up (possibly many) instances, populating them with databases and software, starting the searches, checking the status of the searches, downloading the results, and then shutting down the instances. Accomplishing these tasks requires the user to answer questions such as what instance type is suitable for the BLAST search, whether to use an SSD or some less expensive disk, and where to save the results.

We present ElasticBLAST, a cloud native package that leverages the command-line BLAST+ package to run BLAST on the cloud. There are several reasons to use ElasticBLAST. First, it automates setting up and tearing down instances for the BLAST search, which hides much of the complexity of the cloud and lowers the barrier to entry. At the same time, it makes use of cloud technology where appropriate, leading to a more reliable experience. Second, it can handle anywhere between a few and millions of queries reliably. Third, it distributes the searches to as many instances as the user requests, accelerating the work. Finally, it is supported on AWS and GCP. Given the maturity of cloud technology and the decreasing cost of sequencing (and growing numbers of sequences), ElasticBLAST provides an ideal way to perform alignments in the cloud.

In this article, we describe how ElasticBLAST works, present its user interface, illustrate its use for identifying RNA contamination, and discuss how ElasticBLAST minimizes costs, including using discounted instances, known as spot instances (at AWS) or preemptible instances (at GCP).

## Implementation

### Overall architecture and data flow

ElasticBLAST is a cloud-native, distributed system that runs on GCP and AWS. Its primary aim is to facilitate scalability and ease of use. To this end, ElasticBLAST uses the Platform as a Service (PaaS) [13] cloud service model to facilitate resource management. ElasticBLAST leverages cloud technologies such as container orchestration (Kubernetes), batch compute frameworks (AWS Batch), serverless computing services (AWS Lambda), object storage (also known as buckets, provided by S3 and GCS) as well as block storage devices.

Figure 1 is a schematic of the overall architecture in a cloud service provider agnostic form. There are four primary software modules: query splitting, job submission, resource management, and BLAST. The query splitting and job submission modules can run either on the local machine or as remote jobs on the cloud. The resource manager module runs on the local machine, but some of its functionality can optionally run as a remote job on the cloud if the automatic shutdown feature is enabled. The BLAST module runs only on the cloud.

**Figure 1:**
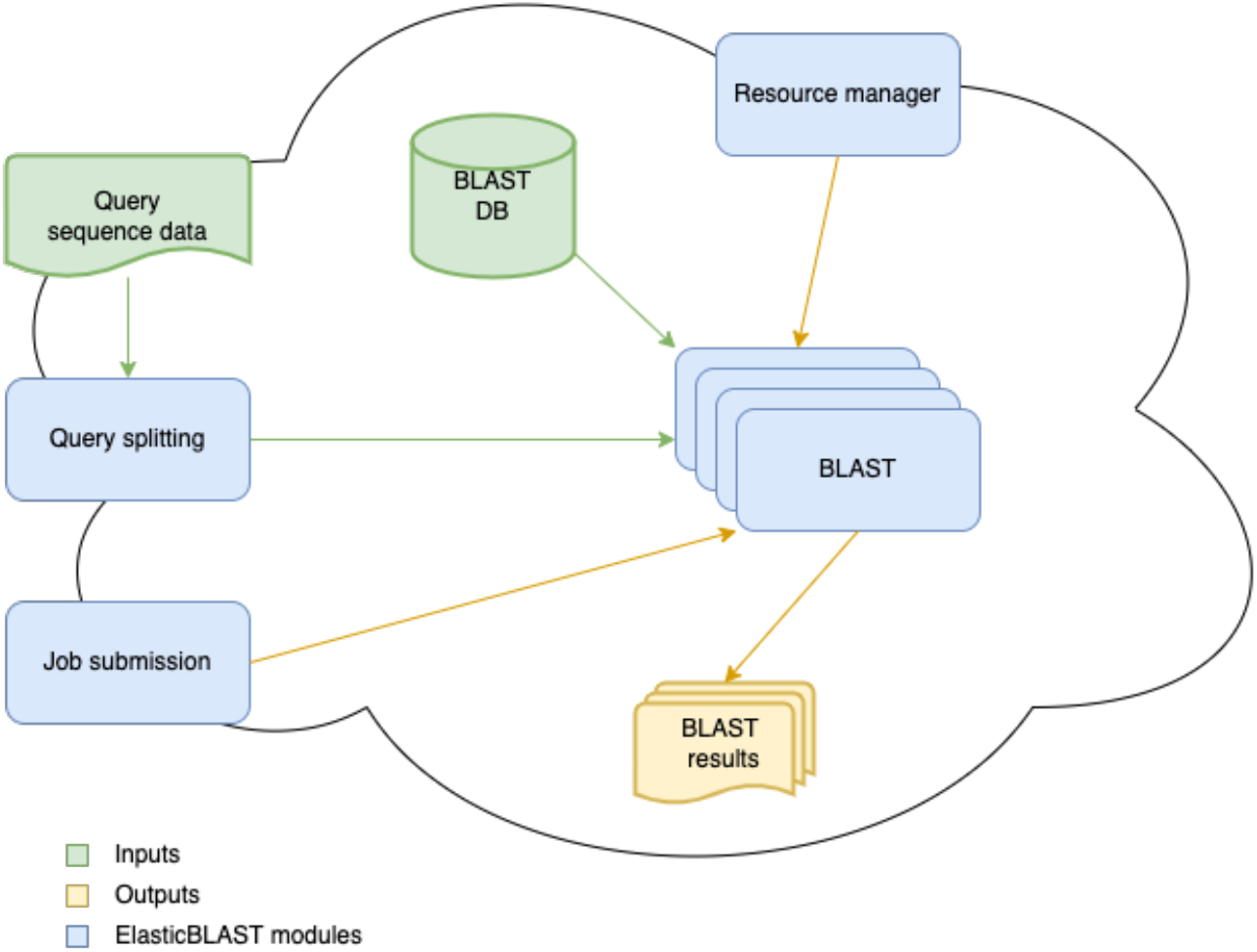
High level ElasticBLAST schematic

The query sequence data can reside on the cloud or on the local machine. It is processed by the query splitting module, which splits it into batches and saves these in cloud storage. Each of the query batches, the choice of BLAST database, and any BLAST parameters specified by the end user constitutes a job, which the job submission module sends to the appropriate managed service for processing. The BLAST module is responsible for providing the query batches to the BLAST+ software [2] for comparison against a BLAST database, which must reside in cloud object storage. The BLAST module can access NCBI-maintained BLAST databases [5] already stored in the cloud as well as custom BLAST databases built and uploaded to the cloud by the end user. The BLAST module saves the result of each of its jobs into the cloud object storage specified by the end user. Figure 2 shows a schematic of the data flow in ElasticBLAST.

**Figure 2:**
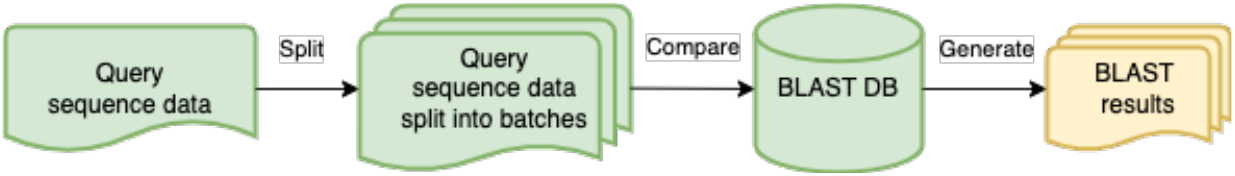
Dataflow in ElasticBLAST

The resource manager module runs on the local machine to allocate cloud resources through a managed service supported by the cloud service provider. It also configures cloud resources to monitor and delete the cloud resources it allocated. The end user can still manually delete cloud resources.

The end user interacts with ElasticBLAST via a command-line application implemented in Python (henceforth referenced as the client or `elastic-blast`). There are three primary subcommands supported by `elastic-blast`: `submit`, `status` and `delete`. The `submit` subcommand processes the ElasticBLAST configuration, creates resources on behalf of the end user and orchestrates query splitting and job submission. The `status` subcommand monitors the ElasticBLAST execution and reports on its progress. The `delete` subcommand initiates the shutdown and deletion of cloud resources allocated by the `submit` subcommand.

Among the critical issues in the ElasticBLAST development was the provisioning, management and efficient use of cloud resources, job orchestration, and query splitting.

### Cluster construction and queuing with cloud native infrastructure

ElasticBLAST relies on managed services for cloud resource management and job orchestration: Google Kubernetes Engine (GKE) [14] in GCP and AWS Batch [15] in AWS.

When configured to run on AWS, ElasticBLAST uses AWS CloudFormation [16], an infrastructure-as-code service that creates “stacks” of resources built from a “template” (Fig. 3, step 1). After the cluster resources are created, the client interacts with AWS Batch (Fig. 3, step 2) to submit jobs to the compute resources allocated (Fig. 3, step 3). The jobs are queued and scheduled to run on instances provisioned by AWS Batch (Fig. 3, step 4). Once the jobs start their execution, they retrieve BLAST databases and query batches from cloud buckets to block storage, and ultimately save the results onto the user’s results bucket (Fig. 3, step 5). AWS Batch handles job failures by retrying them up to three times, unless the jobs ran out of memory, in which case they are flagged as failed. After all BLAST jobs have completed successfully or if any job fails, the resource manager module deallocates the compute resources either through an AWS Lambda function or via the user’s `elastic-blast delete` invocation (Fig. 3, step 6).

**Figure 3:**
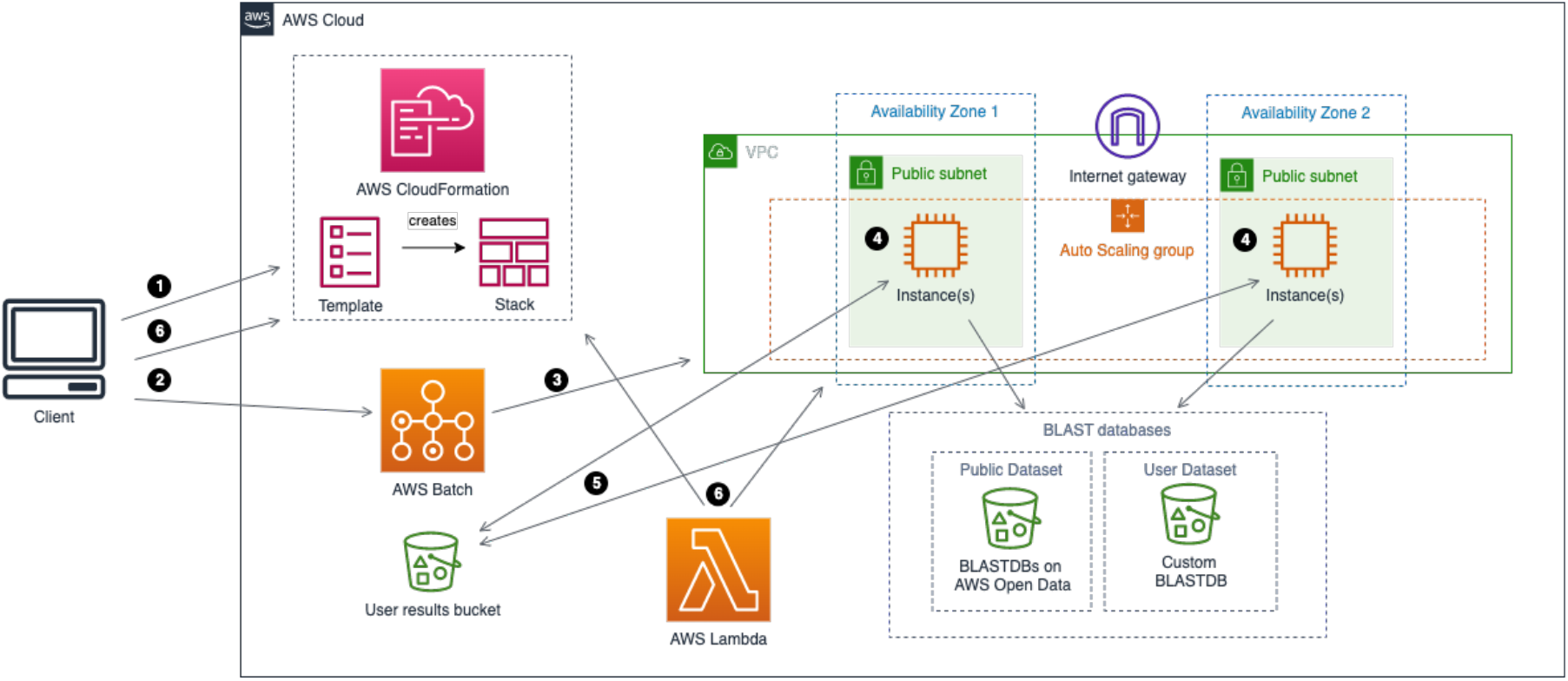
Architecture and workflow overview on AWS

When configured to run on GCP, ElasticBLAST relies on GKE to create a Kubernetes cluster (Fig. 4, step 1). The Kubernetes cluster is configured with a shared persistent disk and an initialization job to retrieve BLAST input data (Fig. 4, steps 2 and 3) onto said disk and split queries. After this initialization completes, the client’s job submission module sends jobs to Kubernetes (Fig. 4, steps 4 & 5) to run BLAST. Kubernetes job objects provide native support for work queues [17], which handle scheduling, queuing, error handling and monitoring of jobs consistent with how it is done in AWS. After all BLAST jobs have completed successfully, their results are written to the user’s cloud bucket (Fig. 4, step 6), and the resource manager module deallocates the compute resources through a Kubernetes cronjob or via the user’s `elastic-blast delete` invocation (Fig. 4, step 7).

**Figure 4:**
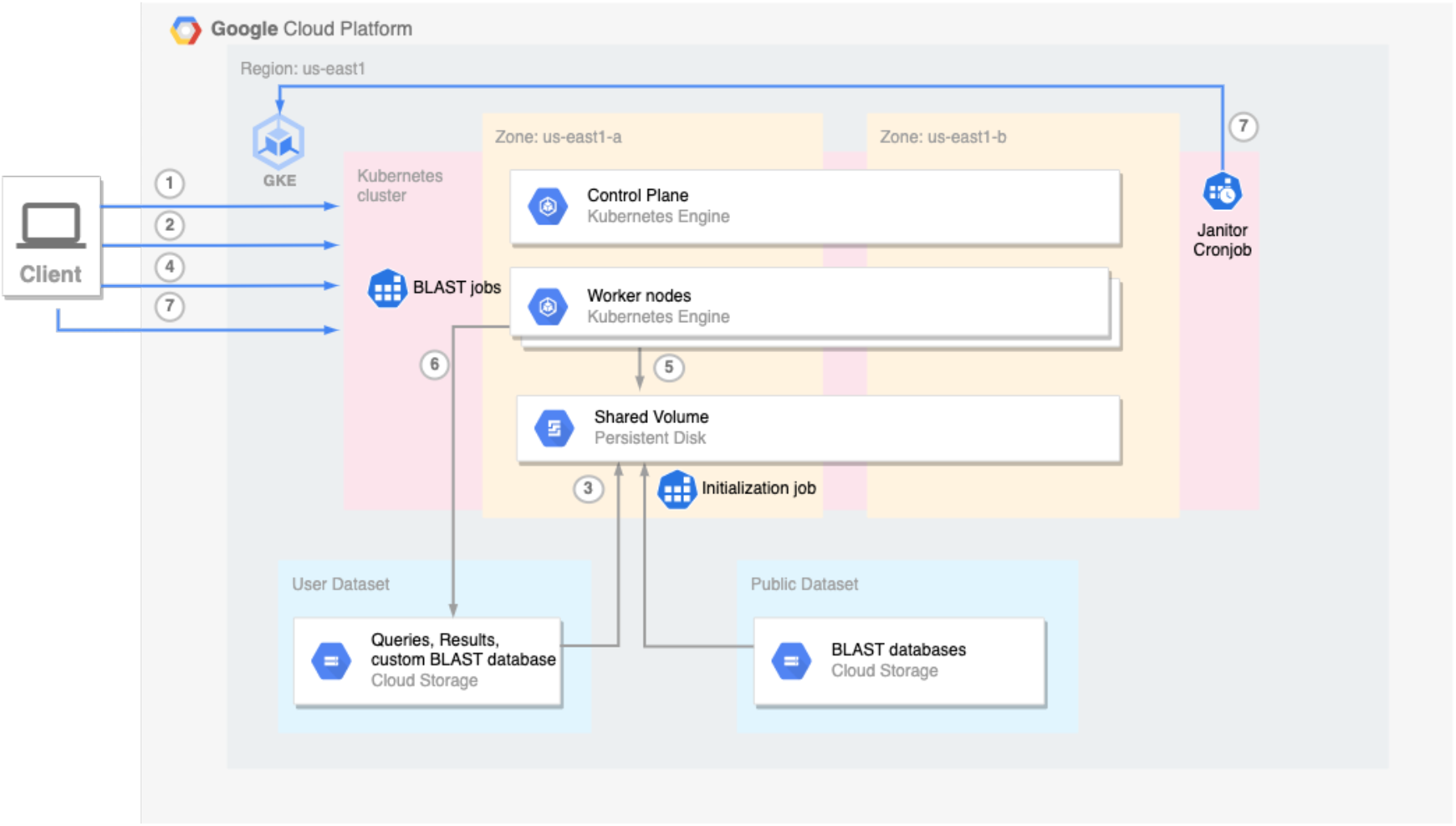
Architecture and workflow overview on GCP

In order to make efficient use of computing resources, ElasticBLAST leverages the horizontal scaling features of GKE and AWS Batch: both managed services will start up as many instances as needed to process the work queue, up to the limit configured by the end user. As the number of outstanding jobs in the queue diminishes, the managed service will shut down the instances when they are no longer needed.

### Query splitting and batch size

ElasticBLAST leverages the cloud to provide multiple worker nodes to parallelize the computation by breaking the queries into query batches. One of the ElasticBLAST parameters that is critical to its performance is the batch length, which specifies the number of bases or residues per query batch. This parameter needs to be configured to a value large enough to amortize the runtime cost of the scheduling and queueing. The authors ran experiments with varying programs, BLAST database sizes, query sizes and batch lengths to arrive at a reasonable default batch length for a given configuration. The goal was to target a median runtime for each of the BLAST jobs in the 5-30 minute time frame. Using this information, ElasticBLAST provides default values for the different BLAST programs, database sizes and query sizes, but the end user can customize this via the `batch-len` [18] configuration parameter.

### Selection of instance types in ElasticBLAST

The choice of instance type is critical to the performance of ElasticBLAST. The instance type provides an upper bound on the number of CPUs and main memory (i.e.: RAM) available to BLAST+. For BLAST+ to run well on a given instance type, the BLAST database must fit in the machine’s main memory. ElasticBLAST relies on BLAST database metadata that is automatically generated when creating BLAST databases (using BLAST+ 2.13.0 or newer) to determine the amount of main memory needed for that database. This in turn is used to select an appropriate instance type from the 400 plus instance types available at each of AWS [19] or GCP [20].

### Automatic shutdown feature

The resource manager module supports an automatic shutdown feature which consists of an AWS Lambda function that monitors the AWS Batch job queue created by ElasticBLAST. The lambda function runs every 5 minutes to check the status of the BLAST jobs and shuts down and deletes all cloud resources, including itself on successful completion of all jobs or the occurrence of a failure. When running in GCP, the resource manager module starts a Kubernetes cronjob to perform the same role as the lambda function in AWS.

## RESULTS and DISCUSSION

We discuss how to run ElasticBLAST and present two search examples, demonstrating its value to the scientific community. In the first case, we show how ElasticBLAST can be used to identify RNA-seq contamination. In the second, we examine how efficiently ElasticBLAST makes use of multiple cloud instances. Finally, we discuss ElasticBLAST in the context of other available tools.

### Interface

A user starts an ElasticBLAST search by invoking the `elastic-blast` application which reads a configuration file specifying the search. Figure 5 shows an example configuration file. There are three sections (cloud-provider, cluster, and blast) that require corresponding information. At the end of the search, the BLAST results are copied to a cloud bucket (owned by the user) specified in the configuration file. The ElasticBLAST documentation [21] provides information on the necessary fields so we will not go into more detail here. It is also possible to use command-line options rather than a configuration file when calling the application, which is also discussed in the documentation.

**Figure 5:**
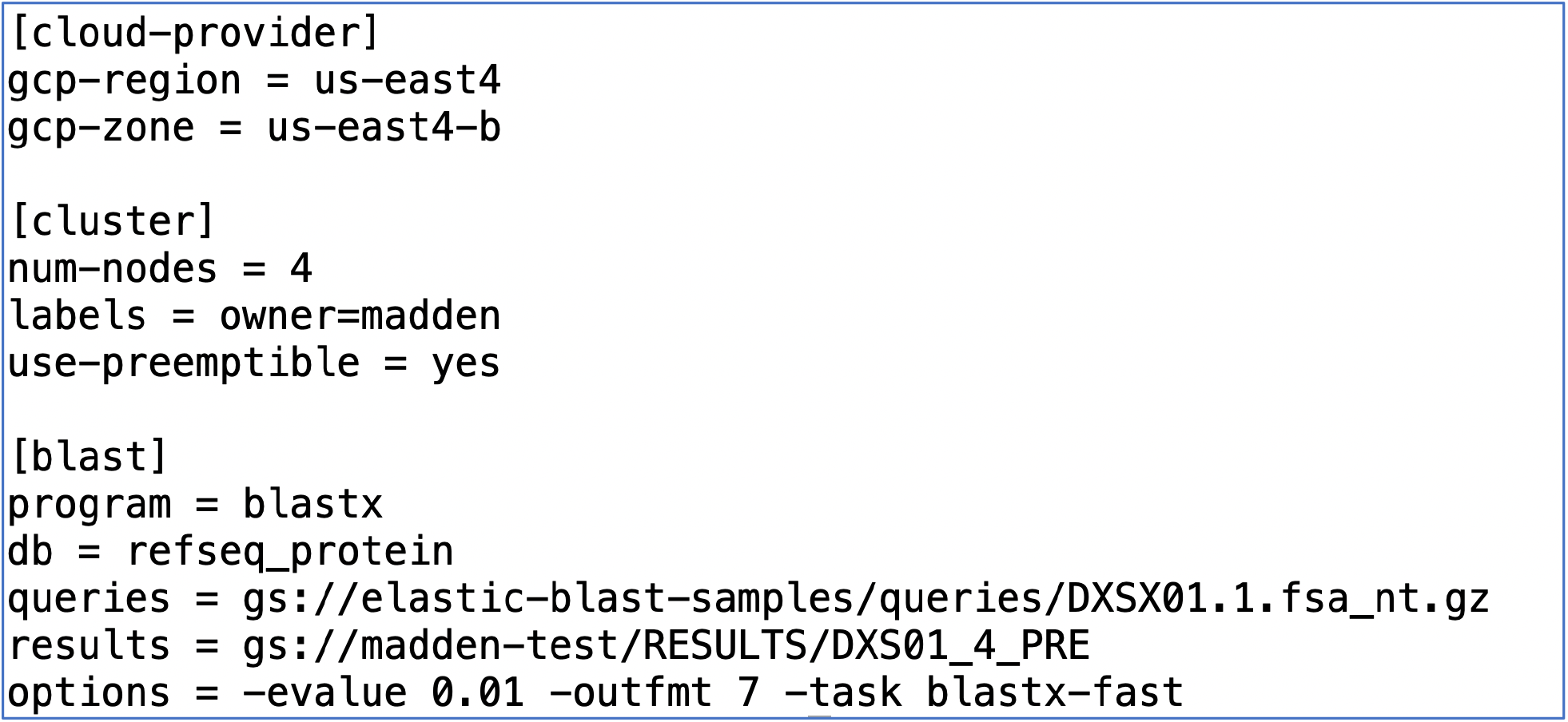
A configuration file used in the second example (below). This configuration file is for GCP. The use-preemptible keyword in the cluster section specifies the use of discounted instances. Information relevant to the search is in the blast section. Results are placed in the user’s bucket specified by the results keyword in the blast section.

As described in the implementation section, ElasticBLAST automatically selects an appropriate instance type for a search, based on database metadata and the BLAST program. The user can override this selection by explicitly setting it in the configuration file.

The ElasticBLAST command-line application can be used as the basis for other interfaces. For example, we have built a Jupyter notebook that calls the elastic-blast application as part of a workflow [22]. It would also be possible to call it based on input to a web page or to containerize the application for use in a pipeline with a formal workflow language.

### Identifying RNA contamination with ElasticBLAST

Whole-transcriptome sequencing (WTS), also known as RNA sequencing (RNA-Seq), is a costeffective means [23] to study differential gene expression profiles [24, 25], phylogenomics [26, 27] or evolution [28, 29]. However, RNA-Seq data analysis is especially challenging if there is no reference genome available in the public databases for the target organism. In this case, a suitable reference transcriptome can be assembled *de novo* and used for quantifying RNA abundance [30].

RNA-Seq contamination, however, is a recognized problem that has played an important role in misleading multiple research conclusions [31, 32]. Detecting and removing contamination prior to a *de novo* transcriptome assembly is a critical step. Nonetheless, detecting contamination in RNA-Seq data is complex due to the sequence similarity between genes in distant taxonomic species. BLAST tools [1, 2] can be used to align RNA-Seq reads to public databases of sequences associating reads with one or more taxonomies. These associations can be used to filter contaminant reads prior to the assembly process. Unfortunately, traditional BLAST searches are time and resource intensive, therefore, k-mer-based methods have been developed to accelerate the computation like Kraken 2 [33]. The improved computing time is at the cost of reducing the sensitivity of the sequence alignments. Although k-mer-based tools are reported to be much faster than programs like BLAST that produce alignments, the latter are still the more sensitive tool for sequence similarity identification [34].

Elastic-BLAST offers a cloud-based solution to efficiently execute BLAST searches in the cloud. The improvement in processing time makes BLAST a usable tool for taxonomic classification of RNA-Seq reads without reducing the sensitivity of the sequence alignments. The GTax database [35] is a taxonomically structured database of genomic sequences comprising 19 taxonomic groups. This database includes representative genome sequences and latest sequence assemblies (if the reference genome is not available) for 20,657 taxa. BLAST searches against GTax are used for taxonomic classification. In the case of an unannotated organism, RNA-Seq reads are aligned to the phylogenetically closest species or remain unidentified after not aligning to any GTax taxonomic group.

We used ElasticBLAST to perform BLAST nucleotide searches (using the blastn executable from the BLAST+ package) against the GTax database to taxonomically classify reads from eight RNA-Seq raw samples from *Physalis peruviana* (TaxID: 126903, BioProject PRJNA67621). This plant is in the *Solanoideae* subfamily (TaxID: 424551) and it is phylogenetically close to the Capsicum genus (peppers, TaxID: 4071) and the *Solanum* genus (TaxID: 4107), which includes flowering plants like tomato, potato, and eggplant.

Our workflow first processes the samples with Trimmomatic [36] to remove adapters and low-quality reads. A total of 26,724,497 reads (5,375,728,710 bases) were aligned with ElasticBLAST against the GTax Eudicotyledons taxonomy group. Figure 6a shows the taxonomy tree created from the BLASTN results with the percentage of reads assigned to each species. The results show that 85.61% of the reads are assigned to species below the correct Pentapetalae clade (TaxID: 1437201). Moreover, 76.85 % of the reads are assigned to species under the correct family Solanaceae (TaxID: 4070). The remaining 14.39% of the reads were aligned to the rest of 18 GTax taxonomic groups. We identified 2.95% of the total reads as contaminants in these samples, see Figure 6b, where the pie chart shows the percentage of reads with respect to total contaminant reads. As expected, the most prevalent contaminants in these samples are the bacteriophage Escherichia virus phiX174 (TaxID: 10847), the fungus Fusarium oxysporum f. sp. lycopersici 4287 (TaxID: 426428) and the bacterium Staphylococcus sp. MZ1 (TaxID: 2836369).

**Figure 6:**
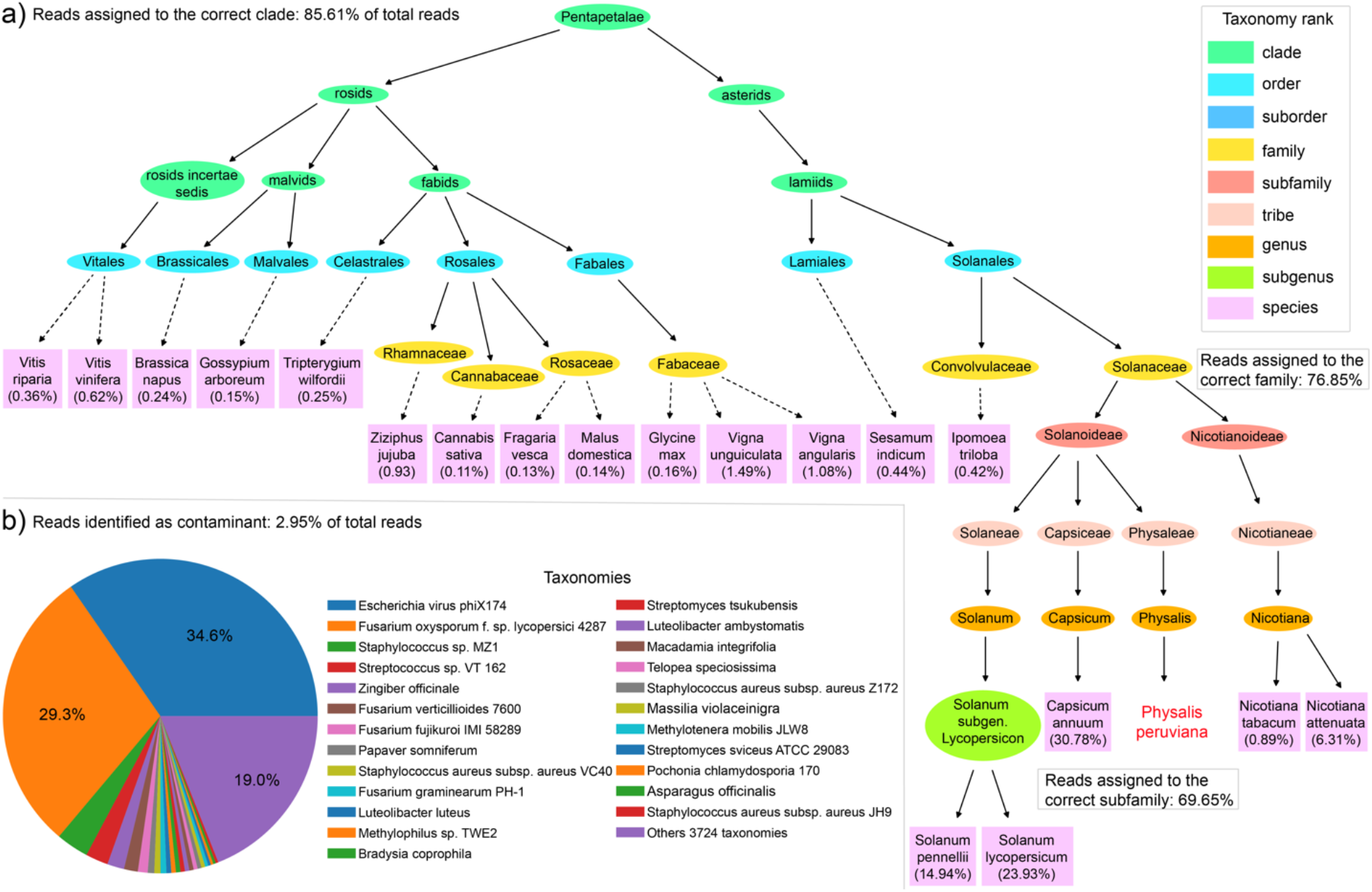
Percent of RNA-Seq reads assigned to each taxonomy species for eight Physalis peruviana samples. a) Taxonomy tree created from the alignment to GTax Eudicotyledons taxonomy group. Percent of reads at species level with respect to total reads in all samples. b) percent of reads not identified in the first alignment that match other GTax taxonomy groups. Percent of reads in the Pie chart are related to the total contaminant reads.

Table 1 presents information on the size of the eudicotyledons and non-eudicotyledons search sets. Table 2 presents information on the cost, run-time, and number of vCPU’s used for the searches. The blastn executable supports multiple search modes, and the sensitive BLASTN mode was used for the taxonomic identification described here. In this example, ElasticBLAST used thousands of instances (44,288 vCPUs) to perform the BLASTN searches of 26,724,497 reads in less than a day at a cost of $0.0013 per read.

**Table 2:** Cost, run-time, and number of virtual CPUs for the ElasticBLAST searches used for taxonomic identification. The costs were calculated by considering the run-time, the hourly on-demand cost per instance, and the number of instances used. Non-eudicotyledons searches were run on an r5ad.4xlarge instance (16 vCPUs, 128 GB RAM, 600 GB local SSD, 10 Gigabit NIC, $1.048/hr on demand). The eudicotyledons searches were run on an c5ad.4xlarge instance (16 vCPUs, 32GB RAM, 600 GB local SSD, 10 Gigabit NIC, $0.688/hr on demand).

| Search set | MegaBLAST time | MegaBLAST cost | MegaBLAST vCPUs | BLASTN time | BLASTN cost | BLASTN vCPUs |
| --- | --- | --- | --- | --- | --- | --- |
| eudicotyledons | 18 minutes | \$20.64 | 1,600 | 3 hours 29 minutes | \$1,849 | 12,288 |
| non-eudicotyledons | 1 hour 19 minutes | \$617.2 | 7,248 | 15 hours 45 minutes | \$33,012 | 32,000 |

We also present information for the megaBLAST search mode, which is optimized for more similar sequences than BLASTN but has a much shorter run-time and is correspondingly less expensive to run. It is suitable for comparing sequences from the same or closely related organisms. The megaBLAST runs cost $0.000024 per read. ElasticBLAST could also start thousands of instances and finish searching 26,724,497 reads in a little more than an hour.

For both megaBLAST and BLASTN an expect value of 0.00001 was used and alignments below 75% identity were discarded. The actual BLAST options were the same in both searches:

`-outfmt “6 qseqid sseqid pident slen length mismatch gapopen qlen qstart qend sstart send evalue bitscore score staxid” - evalue 1e-5 -perc_identity 75 -max_target_seqs 5 -max_hsps 10 - penalty -3`

### Using ElasticBLAST with Multiple Instances

To demonstrate the ability of ElasticBLAST to effectively utilize multiple instances, we present results for ElasticBLAST runs with 1, 2, 4, and 8 instances at GCP. We run these searches with both on-demand and preemptible instances to provide runtime and cost estimates. To provide a baseline, we run BLAST+ on a stand-alone GCP instance using a script to execute operations performed by ElasticBLAST to configure and run BLAST. For this series of runs we searched 224 contigs (631,309 nucleotides) from the WGS project (DXSX00000000.1) for *Candidatus Saccharibacteria bacterium* against the refseq_protein database, consisting of high-quality proteins from the NCBI RefSeq project [37]. We used the BLASTX program, which translates a nucleotide query in six frames and compares it to a protein database. ElasticBLAST selected the e2-highmem-16 instance with sufficient memory (128 GB) to accommodate the database (90 billion residues). Table 3 presents the results, and we discuss those below. The BLAST parameters used for these searches were:

~~~
-outfmt 7 -evalue 0.01 -task blastx-fast
~~~

**Table 3:** ElasticBLAST searches of *Candidatus Saccharibacteria bacterium* contigs from the DXSX00000000.1 project against the refseq_protein database. Searches were performed at GCP with the e2-highmem-16 instance, selected by ElasticBLAST. The cost was estimated using the search time, the number of instances, and the hourly cost ($0.81432) of the instances (as of October 25, 2022). Costs for ElasticBLAST runs on preemptible instances were calculated at 20% of on-demand instance cost and are shown in the last two columns. The last row (“standalone”) shows data for a stand-alone BLAST+ run, performed on one on-demand instance.

| Number of instances | Time (hours) | Cost (\$) | Time (hours) preemptible | Cost (\$) preemptible |
| --- | --- | --- | --- | --- |
| 1 | 8.07 | 6.57 | 7.83 | 1.28 |
| 2 | 4.08 | 6.44 | 4.13 | 1.30 |
| 4 | 2.37 | 6.87 | 2.22 | 1.28 |
| 8 | 1.63 | 8.55 | 1.5 | 1.59 |
| 1 (stand-alone) | 6.62 | 5.39 | NA | NA |

Discounted instances are an effective means to reduce cloud costs. At GCP, they are called preemptible instances and cost 20% of the on-demand instance price. ElasticBLAST searched the 631,309 nucleotides against the 90 million residues in refseq_protein for a little more than a dollar with one instance, but this run costs about $6 with one on-demand instance (first row of Table 3). We did not have a problem acquiring discounted instances for our runs, and they ran in roughly the same time (and sometimes less) as the on-demand instances.

ElasticBLAST can effectively use multiple instances. The run-time with two instances was about 50% of that for a single instance, and the cost was about the same. The run-time with four instances was about 29% of the single instance time, and the cost was 5% higher. The search with eight instances cost about 30% more than the single instance run and ran in about 20% of the single instance time. The extra expense of the eight-instance run is due to the time it takes Kubernetes to detect that instances are no longer needed and shut them down (data not shown), but the run is still under $2 using preemptible instances. Figure 7 presents a screenshot of the GCP monitoring view for the four-instance run, consisting of two graphs. The top graph shows the number of instances in the cluster at a given time. The bottom graph shows how busy the cluster is at a given time. Every ElasticBLAST search on GCP has a start-up period where only one instance is running allowing for cluster configuration and BLAST database installation. This time varies, depending upon the state of the network at the provider and the size of the database, but was 22 minutes or less for the runs discussed here. The bottom graph shows that the cluster is no longer completely busy after 4:20, and the top graph shows that cluster shrinking in response, completely shutting down after all instances are no longer busy.

**Figure 7:**
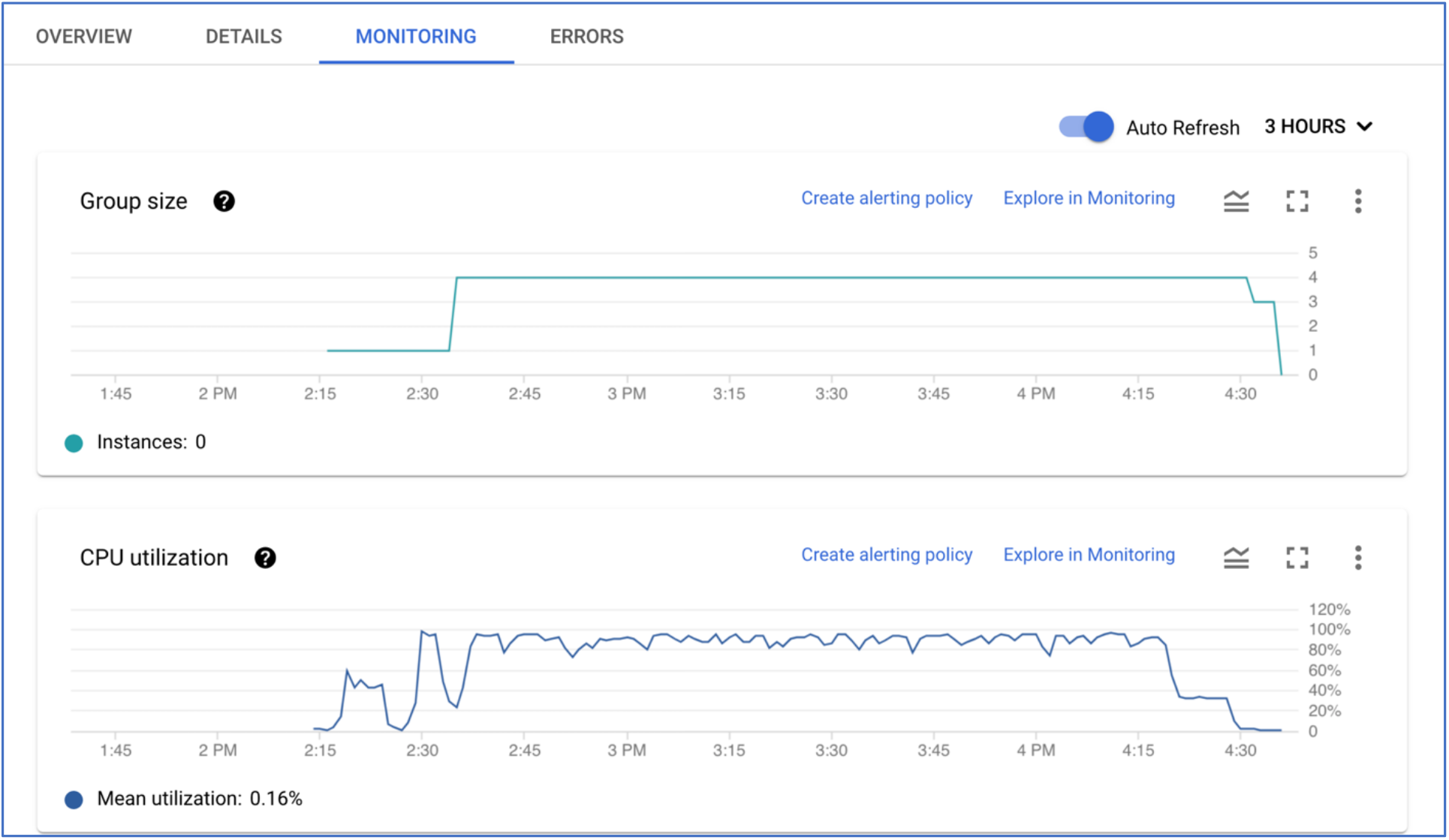
Cluster size (top) and CPU utilization of the cluster (bottom) for an ElasticBLAST run with four instances. This is a screenshot of the GCP monitoring view for the cluster. The cluster has only one instance from 2:15-2:40 (top graph), allowing for the installation of software and databases. The bottom graph shows that the cluster has about 50% CPU utilization after 4:20, and the top graph shows the cluster size shrinking about 10 minutes later. The CPU utilization at a given time is based on the size of the cluster at that point in time.

For the stand-alone run, a bash script was used to download the BLAST+ package, the refseq_protein BLAST database and the query file, and then start the BLAST+ search (using 15 threads). The time and cost shown in Table 3 include all those tasks. The stand-alone run was faster than ElasticBLAST with a single instance but did not include starting the instance, downloading results to a cloud bucket (after the search) and shutting the instance down.

ElasticBLAST is most efficient working on large query sets. There are a few reasons for this. First, each ElasticBLAST search involves setup overhead, which occurs once regardless of the number of queries. Second, ElasticBLAST groups the queries into batches that are large enough to amortize the scheduling overhead of any individual batch. In the example above the setup time is around 20 minutes, most of which is to download the refseq_protein database (135 GB).

We have discussed searches with a wide range of costs. A relatively small search of a few hundred contigs against a smaller (but well curated) protein database cost less than $2 with discounted instances. A large search of more than 5 billion bases against a database of more than 400 billion bases using a sensitive search algorithm cost about $33,000. There are some best practices to prevent surprises. First, as noted in [8], it is a good idea to perform a small test run before submitting large jobs. This allows users to estimate overall cost and possibly change strategy. Second, discounted instances can provide substantial savings, and ElasticBLAST makes it straightforward to use those. Cautious users can specify a small number of instances and, if it seems the run will cost too much, stop the searches with the delete command. Already processed results will be available in the cloud bucket.

ElasticBLAST implements several best practices to make searches efficient. First, it selects appropriate hardware for the search. It uses SSDs for the databases, which are more expensive than spinning disks, but inexpensive compared to the cost of the instances and respond more quickly, keeping the databases in memory and the instances busy. It also selects the smallest instance, based on the size of the database, that has sufficient memory to hold the database and allow BLAST to function efficiently. Second, it uses cloud services that shut down instances when they are no longer needed. Third, it optimizes the BLAST threading for the search, using at most 16 threads per process as this is optimal for BLAST. Instances with more than 16 vCPUs (e.g, 32 vCPUs) run multiple processes. For small databases, it uses a threading model that is more efficient for those databases [38].

### Other Tools

We compare ElasticBLAST to other software that can run BLAST on the cloud. We limit our discussion to projects that make source code available and can be run without a licensing fee to match those features of ElasticBLAST.

First, we discuss two packages that use cloud technology to produce sequence alignments. SparkBLAST [39] is a cloud native application that runs BLAST on GCP and Azure. It uses cloud native technology to distribute queries over multiple instances running an unsupported version of BLAST (“blastall”). It does not create or provision instances for the BLAST search and does not support spot instances. SparkBLAST is from 2017 and does not appear to be currently in development. Sparky-BLAST [40] is an application that runs its own implementation of BLAST (written in Python). It is presented by the authors as a proposal, but it has some interesting features. It can distribute a database across multiple instances, allowing Sparky-BLAST to use smaller machines for the searches. ElasticBLAST is unable to distribute databases in this manner, but cloud providers offer machines with enough memory to accommodate most BLAST databases. Sparky-BLAST can also distribute a set of queries over multiple instances, and the authors demonstrate that it scales well (with five 16 CPU instances) and compare it to BLAST+ running on one instance, since BLAST+ cannot use multiple instances. ElasticBLAST, running BLAST+ on the cloud for the user, lifts this limitation by distributing searches over multiple instances. As we have shown above, it also can scale the searches with different numbers of instances. Sparky-BLAST only supports one BLAST program (BLASTN or DNA-DNA comparisons), requires the user to set up a SPARK cluster and Cassandra database, and does not offer the full range of BLAST report options so does not seem suitable for most uses of BLAST. The authors in [40] also do not make clear which cloud provider was used for their benchmarking or if it was run on non-cloud machines. It does make interesting and innovative use of cloud technology to improve sequence searches.

Nextflow [41] is a general package for running a pipeline, and one could use it to run BLAST searches. It supports spot instances, has cloud support, and can create and provision instances, but the user must provide the software. It is unable to decide which instances will be able to run a BLAST search, unlike ElasticBLAST. It is specifically targeted to “bioinformaticians familiar with programming” [41], whereas ElasticBLAST does not require programming experience.

ElasticBLAST was designed specifically for the cloud, with the goal of making it easy to run there. There is other software that will run BLAST on the cloud, but nothing with the functionality of ElasticBLAST.

## CONCLUSION

We presented ElasticBLAST, a new cloud native application that can run BLAST+ searches on a cloud provider. ElasticBLAST simplifies running a BLAST search on the cloud. It can choose a cloud instance suitable for a BLAST search, given information about the database and program. It can also use discounted instances, saving the user money. ElasticBLAST can search NCBI or user provided databases and supports most of the BLAST+ programs and options. It is supported at both AWS and GCP.

Extensive documentation for ElasticBLAST is available at [21]. This documentation includes an introduction to the cloud and ElasticBLAST as well as quickstarts for GCP and AWS, so a researcher can try out ElasticBLAST with minimal effort. Tutorials and documentation for parameters are also available. The ElasticBLAST source code is available at GitHub [42]. We also provide a GitHub repository with scripts that use ElasticBLAST [22], which includes a Jupyter notebook.

We are exploring ways to improve ElasticBLAST. These include optimizing the setting of parameters (e.g., batch-len) used by ElasticBLAST and improving the ability of ElasticBLAST to read in large numbers of sequences from SRA. We are also interested in integrating ElasticBLAST into workflows. We welcome feedback from users on features that would make ElasticBLAST more useful.

## Availability and Requirements

Project Name: ElasticBLAST

Project Home Page: https://github.com/ncbi/elastic-blast

Operating Systems: 64-bit LINUX, MacOS

Programming Language: Python

Other Requirements: See https://blast.ncbi.nlm.nih.gov/doc/elastic-blast/requirements.html

License: Public Domain [43]

Any restrictions to use by non-academic users: none

## Abbreviations

AWS: Amazon Web Services
BLAST: Basic Local Alignment Search Tool
GCP: Google Cloud Platform
GCS: Google Cloud Storage
GKE: Google Kubernetes Engine
PaaS: Platform as a Service
RAM: Random Access Memory
RNA-seq: RNA sequencing
SSD: Solid State Drive
S3: Simple Storage Service
vCPU: virtual central processing unit
WTS: Whole-transcriptome sequencing

## Declarations

### Ethics approval and consent to participate

Not applicable

### Consent for publication

Not applicable

### Availability of Data and Materials

The software is freely available on GitHub at https://github.com/ncbi/elastic-blast All datasets analyzed during this study are freely available. The SRA runs used for the first example were SRR1944534, SRR1945431, SRR1952996, SRR1955167, SRR1955548, SRR1955886, SRR1957684, and SRR1958937

### Author Contributions

CC, GB, TM conceived the project. CC, GB, VJ implemented the project. TM and RV ran experiments for the paper. CC, GB, RV, and TM drafted the manuscript. TM directed the project.

## Acknowledgements

We would like to thank Vadim Zalunin, Anatoliy Kuznetsov, and Eugene Yaschenko for valuable architectural discussions. We would like to thank Scott McGinnis, Preye Akuiyibo, Dave Arndt, Priyanka Ghosh, Tao Tao, Peter Cooper, Wayne Matten, Rich McVeigh, Ryan Connor, Ravinder Eskandary, Yan Raytselis, and Yuri Merezhuk for useful discussions, feedback, and help. We would like to thank Kim Pruitt, Valerie Schneider, Wratko Hlavina, Rodney Brister, Bart Trawick, and David Landsman for their support.

